# Pseudouridines in the U2 Branch Site Recognition Region Differentially Contribute to Branch Site Recognition during Pre-mRNA Splicing

**DOI:** 10.64898/2026.08.09.743773

**Authors:** Rui Zhang, Meemanage D. De Zoysa, Jonathan L. Chen, Hironori Adachi, Yu Sun, Yi-Tao Yu

**Author notes:** Corresponding author: Yi-Tao Yu, Department of Biochemistry and Biophysics Center for RNA Biology University of Rochester Medical Center Box 712 601 Elmwood Avenue Rochester, NY14642 USA.

## Abstract

Pseudouridines (Ψs) are highly enriched in conserved, functionally critical regions of spliceosomal snRNAs, particularly within the U2 branch site recognition region (BSRR), which contains six Ψs in humans and three in *S. cerevisiae*. To investigate how U2 BSRR pseudouridylation influences branch site sequence (BSS) recognition, we developed a large-scale, high-throughput screening system in *S. cerevisiae* that allows a library of pre-mRNAs with randomized BSSs to be spliced in distinct U2 BSRR pseudouridylation backgrounds. Screening and next-generation sequencing (NGS) revealed that different U2 pseudouridylation backgrounds exhibit distinct recognition patterns and efficiencies for specific BSSs. Notably, Ψ42 and Ψ44 generally enhanced splicing, whereas Ψ38 alone, and in some contexts together with Ψ35, consistently impaired BSS recognition. The differential effects were validated using endogenous *S. cerevisiae* genes. In addition, splicing assays with engineered pre-mRNA constructs guided by the screening results demonstrated that BSRR Ψs directly influence BSS selection, supporting a model in which U2 pseudouridylation modulates BSS recognition and could contribute to alternative splicing in more complex eukaryotes. Finally, synthetic-lethality analyses with a Prp5 mutant, together with Prp5-U2-pre-mRNA binding assays, indicate that U2 BSRR Ψs are critical for Prp5 recruitment and, consequently, for proper U2-BSS interactions during spliceosome assembly. Collectively, these findings establish U2 pseudouridylation as a key determinant of branch site recognition and spliceosome function.

## Introduction

Pre-mRNA splicing, the process of removing introns from pre-messenger RNA, is a fundamental and tightly regulated mechanism of gene expression in eukaryotes (Wahl et al. 2009; Lee and Rio 2015; Wilkinson et al. 2020). This process is carried out by the spliceosome, a large and dynamic RNA-protein complex consisting of hundreds of proteins and five small nuclear RNAs (snRNAs): U1, U2, U4, U5, and U6. These snRNAs themselves exist in the cell as RNA-protein complexes known as small nuclear ribonucleoproteins (snRNPs), which directly participate in spliceosome assembly (Wahl et al. 2009; Wilkinson et al. 2020).

Spliceosome assembly occurs in a stepwise manner (Moore et al. 1993; Staley and Guthrie 1998; Yu et al. 1999; Jurica and Moore 2003; Fabrizio et al. 2009; Will and Lührmann 2011; Hoskins and Moore 2012). The process begins when the U1 snRNP recognizes and binds to the 5’ splice site of the pre-mRNA. U2 auxiliary factor (U2AF) and splicing factor 1 (SF1) then recognize the 3’ splice site and the branch site, respectively, facilitating the recruitment of the U2 snRNP to the branch site sequence (BSS) (Parker et al. 1987; Zhuang and Weiner 1989). Concurrently, the DEAD-box ATPase Prp5 is recruited through binding to the U2 branchpoint-interacting stem loop (BSL), exposing the U2 branch site recognition sequence for pairing with the BSS (Xu and Query 2007; Perriman and Ares 2010; Liang and Cheng 2015; Wu et al. 2016). ATP hydrolysis by Prp5 then helps drive the transition to a productive pre-splicing complex (complex A) by stabilizing correctly formed U2-BSS interactions. Concomitant with Prp5 release, the U4/U6/U5 tri-snRNP is recruited (Liang and Cheng 2015), resulting in formation of the precatalytic spliceosome. Following extensive conformational rearrangements, the spliceosome catalyzes the two transesterification reactions of splicing, resulting in exon ligation and release of the intron as a lariat structure (Wahl et al. 2009). Throughout the process, U2 remains base-paired with the BSS of the pre-mRNA (Zhuang and Weiner 1989; Moore et al. 1993; Yu et al. 1999).

It is well known that all five higher eukaryotic spliceosomal snRNAs are pseudouridylated and that these modifications are generally conserved across species (Reddy and Busch 1988; Gu et al. 1996; Massenet et al. 1999; Yu et al. 1999). Notably, these Ψs are predominantly located in regions involved in RNA-RNA and RNA-protein interaction networks during spliceosome assembly, suggesting their important functional roles in pre-mRNA splicing. Among all spliceosomal snRNAs, U2 snRNA is the most extensively pseudouridylated (Reddy and Busch 1988; Yu et al. 1998; Yu et al. 2011). Despite the fact that the U2 branch site recognition region (BSRR) is identical across species, the number of Ψs within this region varies substantially between yeast and higher eukaryotes. For instance, in vertebrate U2, all six uridines in the BSRR are converted to Ψ (Ψ34, Ψ37, Ψ39, Ψ41, Ψ43, and Ψ44). In contrast, there are only three Ψs in *S. cerevisiae* U2 snRNA (Ψ35, Ψ42, and Ψ44, corresponding to Ψ34, Ψ41, and Ψ43 in vertebrates). Genetic studies in yeast and microinjection experiments in Xenopus oocytes have demonstrated that these U2 BSRR Ψs in combination are critical for splicing (Zhao and Yu 2004b; Yang et al. 2005; Zhao and Yu 2007; Wu et al. 2016). However, how individual Ψs, as well as their different combinations within the U2 BSRR, contribute to BSS recognition and pre-mRNA splicing remains poorly understood.

In this study, we developed a yeast-based screening system to systematically investigate the roles of U2 BSRR Ψs in branch site recognition and pre-mRNA splicing. Our results demonstrate that distinct U2 BSRR pseudouridylation patterns differentially influence branch site recognition and selection. Mechanistically, these Ψs appear to influence the binding of Prp5, which plays an important role in the quality control of U2-BSS interactions during early spliceosome assembly.

## Results

### Developing a new yeast system to screen for favorable U2 BSRS-BSS interactions

To conduct a comprehensive evaluation of the contributions of Ψs to BSS recognition, we developed a *S. cerevisiae* screening system, in which the influence of Ψ on U2 BSRS-BSS recognition can be directly assessed.

We employed the *ACT1-CUP1* reporter system, originally established by the Guthrie group (Lesser and Guthrie 1993), to measure the efficiency of splicing of the *ACT1-CUP1* reporter pre-mRNA. *CUP1* encodes the Cup1 protein, which chelates Cu^2+^ ions and enables cell survival in a copper-containing medium in a dose-dependent manner. Upon transformation with a plasmid harboring the *ACT1-CUP1* fusion reporter gene, which includes a partial ACT1 gene (the first exon, the intron, and a small segment of the second exon) fused to a full-length *CUP1* gene (**Fig 1A**), the *Cup1Δ* strain synthesizes *ACT1-CUP1* pre-mRNA. Splicing of this pre-mRNA yields mature *ACT1-CUP1* mRNA, which is then translated into the functional Act1-Cup1 fusion protein, enabling cell growth in Cu^2+^-containing medium. Previous studies have shown a strong correlation between splicing efficiency and the production of the Act1-Cup1 fusion protein. The amount of fusion protein expressed in cells is directly reflected by cell proliferation in media containing various Cu^2+^ concentrations ([Cu^2+^]) (Lesser and Guthrie 1993; Hilliker et al. 2007; Perriman and Ares 2007; Xu and Query 2007). Therefore, by determining the range of [Cu^2+^] that permits cell growth, one can assess the relative efficiency of splicing for the *ACT1-CUP1* fusion pre-mRNA.

**Fig. 1.**
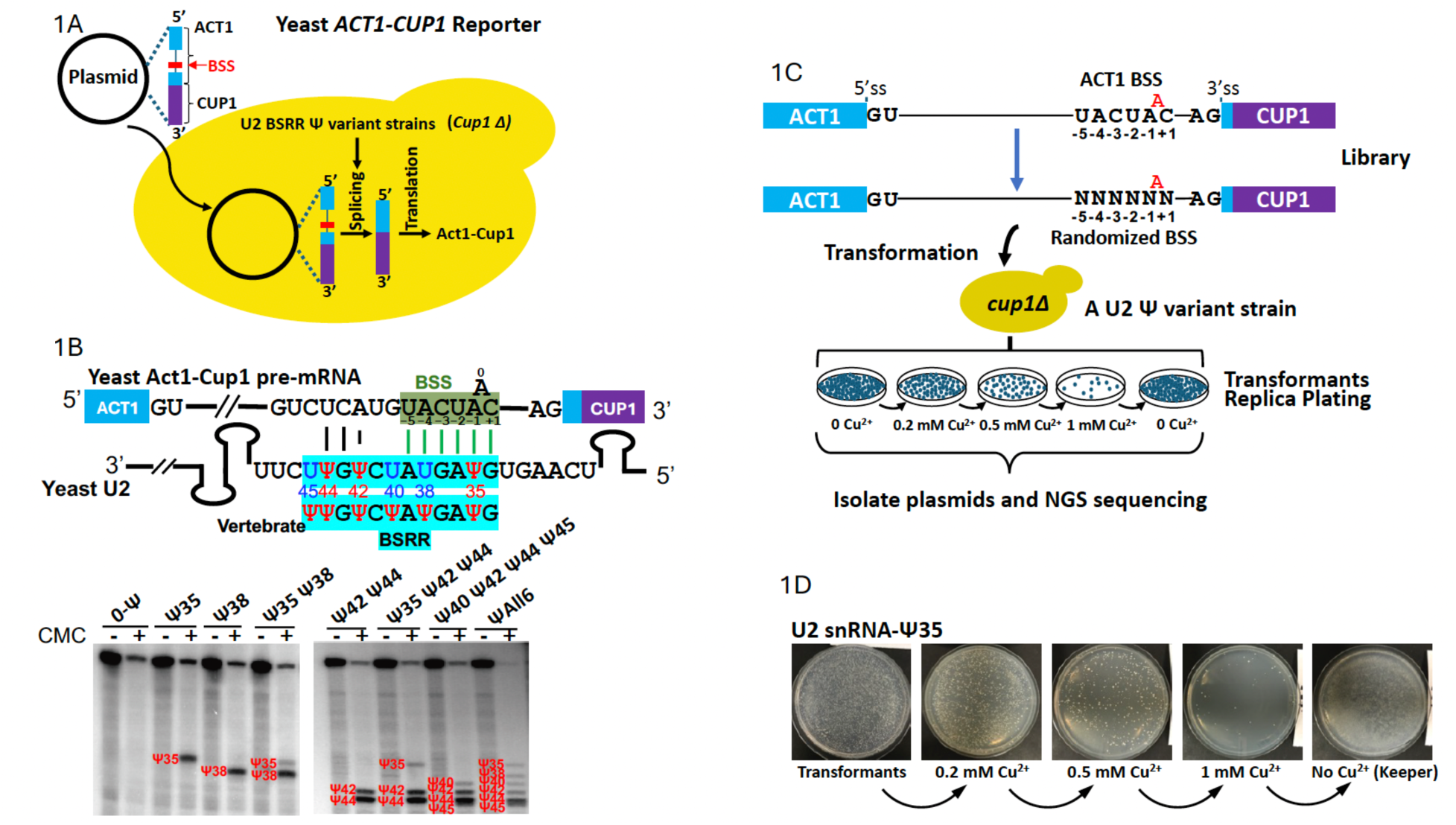
Establishment of a yeast screening system. (A) Schematic of the *ACT1-CUP1* reporter. Yeast *ACT1* containing a truncated exon 2 was fused to the coding sequence of the *CUP1* gene. Following transformation of the reporter into the *cup1Δ* strain, the efficiency of *ACT1-CUP1* pre-mRNA splicing was assessed by monitoring yeast growth on media containing different concentrations of Cu^2+^. (B) Top panel: Diagram of *ACT1-CUP1* pre-mRNA-U2 interactions. The Ψ residues and their positions in the yeast and vertebrate U2 BSRR are indicated. Bottom panel: Detection of U2 Ψ residues by CMC-based primer extension. CMC–, without CMC treatment, yielding a Ψ-readthrough primer-extension product; CMC+, with CMC treatment, resulting in primer-extension stop(s) one nucleotide before the Ψ site. Strains are indicated at the top. The detected signals (bands) and the red Ψ numbers correspond to positions within the yeast BSRR. (C) Schematic of the *ACT1-CUP1* BSS library and the large-scale screening strategy. ‘N’ denotes a randomized nucleotide, and the red ‘A’ indicates the branch point. Library plasmids were transformed into *cup1Δ* strains carrying U2 BSRR Ψ variants. Transformants were replica plated onto agar plates containing various concentrations of Cu^2+^. Colonies were then harvested for plasmid extraction and NGS. (D) Representative example of replica plating using the U2 Ψ35 strain.

To analyze the impact of Ψs in the U2 BSRR on BSS recognition, we created a set of *cup1Δ* yeast strains, each carrying a different U2 variant with a distinct number and combination of Ψs within the BSRR (**Supplementary Table 1**). Specifically, we took advantage of our ability to manipulate pseudouridylation within the U2 BSRR. In the wild-type *S. cerevisiae*, Ψ35, Ψ42, and Ψ44 are modified by Pus7, snR81, and Pus1, respectively; no redundant Ψ synthases act at these positions (Massenet et al. 1999; Ma et al. 2003; Ma et al. 2005; Deryusheva and Gall 2017). While Pus7 and Pus1 are stand-alone protein Ψ synthases, snR81 is a box H/ACA ribonucleoprotein (RNP) consisting of a box H/ACA RNA and four core proteins (Nhp2, Nop10, Gar1, and Cbf5) (Ma et al. 2005; Hamma and Ferré-D’Amaré 2010; Kiss et al. 2010). To generate *cup1Δ* strains with different U2 Ψ variants but otherwise identical genetic background, we first deleted *PUS1*, *PUS7,* and *SNR81*, yielding a “0-Ψ” strain of no Ψs in the U2 BSRR. We then introduced various designer box H/ACA guide RNAs (gRNAs) (De Zoysa and Yu 2017; De Zoysa et al. 2018; Adachi et al. 2023) to direct site-specific pseudouridylation at any of the six uridines within the U2 BSRR, generating *cup1Δ* strains with distinct Ψ numbers and combinations. For screening purposes, we initially created **eight** such *cup1Δ* strains (**Supplementary Table 1**), each containing a different U2 Ψ variant. All strains exhibited normal growth under optimal nutrient conditions (no Cu^2+^), compared to the wild-type strain. To confirm that the designer box H/ACA gRNAs successfully directed pseudouridylation to the intended sites, we performed the 1-Cyclohexyl-3-(2-morpholinoethyl) carbodiimide metho-p-toluenesulfonate (CMC)-modification-based assay (Ofengand et al. 2001; Carlile et al. 2014; Schwartz et al. 2014; Li et al. 2015) using total RNA purified from each strain. These experiments confirmed that the designer gRNAs functioned exactly as expected, with Ψ(s) detected at the precise target sites specified by the corresponding designer gRNA(s) (**Fig 1B**).

We next constructed an *ACT1-CUP1* library, in which the BSS sequence was randomized. Specifically, six of the seven nucleotides (excluding the branch point adenosine, ‘A’) of the *ACT1* BSS were randomized, theoretically yielding 4096 different BSSs (4^6^) (**Fig 1C**). To ensure that the library was constructed correctly, we subjected it to Amplicon-EZ sequencing (short-read, high throughput NGS), and identified 4011 unique BSSs (from more than 15,000 *E.coli* transformants), covering ∼98% of the theoretical diversity (**Supplementary Fig 1A**). The mean read count per BSS was 53.5; notably, the most naturally conserved BSS (UACUAAC) had a read count of 55, closely matching the average and approaching the theoretical ideal. This *ACT1-CUP1* library, in combination with the **eight** U2 Ψ variant *cup1Δ* strains, provided a new screening system for identifying favorable U2 BSRS-BSS interactions.

### High-throughput screening uncovers BSS landscapes under U2 Ψ variant conditions

To analyze the impact of Ψs in the U2 BSRR on BSS recognition and selection, we transformed each of the **eight** U2 Ψ variant *cup1Δ* strains with the *ACT1-CUP1* library, in which the BSS was randomized. After transformation, a large number of single colonies grew out on the agar plates lacking Cu^2+^. These colonies were replica-plated onto agar plates containing two [Cu^2+^] concentrations (0.2 mM and 0.5 mM). As anticipated, more yeast colonies grew on the plates containing low [Cu^2+^] than on those with high [Cu^2+^] (**Fig 1D**). Colonies from each plate were harvested, and plasmid DNAs were extracted from the cells. Upon amplification with barcode primers, the *ACT1-CUP1* gene, which contains various BSSs, was sequenced.

By comparing the BSSs from the 0-[Cu^2+^] plates with the BSSs from the untransformed library (**Supplementary Fig 1B**), we observed a linear correlation between the two; no significant differences in sequence composition or read counts were detected, indicating that the transformation and screening experiments were reliable. The BSS groups obtained from the 0-Cu^2+^ plates were subsequently used to normalize all Cu^2+^-screened datasets. Splicing efficiency was then inferred from the normalized read counts of BSSs recovered from the Cu^2+^-containing plates.

As shown in **Fig 2A**, the BSSs selected from all eight strains under two [Cu²⁺] concentrations (0.2 mM and 0.5 mM) were evaluated after normalizing to the 0-[Cu²⁺] control. As **Fig 2A** shows, more BSSs were selected from the 0.2 mM [Cu²⁺] condition than from the 0.5 mM [Cu²⁺] condition. Among the selected sequences, the canonical branch site sequence UACUA<u>A</u>C (the underlined A is the branchpoint adenosine), which base-pairs perfectly with U2, was one of the most favorably selected for all eight U2 Ψ-variant strains (**Fig 2A**). In addition, all 11 known native yeast BSSs were selected. Notably, as shown in **Fig 2B**, the uridine located two nucleotides upstream of the branchpoint adenosine (U_-2_) was virtually invariant among all selected sequences. Similarly, the nucleotide immediately upstream of the branchpoint adenosine was most frequently adenosine (A_-1_), although guanosine appeared in some instances. Two cytosines, C_-3_ and C_+1_, positioned three nucleotides upstream and one nucleotide downstream of the branchpoint adenosine, respectively, were also relatively conserved, although one or both were sometimes replaced by uridine. Importantly, the conservation levels at these nucleotides were not strictly uniform across all eight strains. In particular, C_+1_ was more highly conserved in the Ψ38 strain than in any of the other strains. Also notable was the fact that although the first two nucleotides of the BSS showed relatively little conservation overall, they exhibited more variability in the Ψ35 Ψ42 Ψ44, Ψ42 Ψ44, and Ψ40 Ψ42 Ψ44 Ψ45 strains than in the 0-Ψ, Ψ38, and Ψ35Ψ38 strains. Consistent with these observations, the fewest BSSs were selected from the Ψ38 and Ψ35Ψ38 strains, whereas substantially more were selected from the Ψ35 Ψ42 Ψ44, Ψ42 Ψ44, and Ψ40 Ψ42 Ψ44 Ψ45 strains. Moreover, Ψ35 alone, or the absence of Ψs in the BSRR (0-Ψ strain), conferred a slight advantage relative to Ψ38, enabling recognition of a slightly broader range of BSSs. Together, these results suggest that Ψ38 in the BSRR imposes more stringent constraints on U2-BSS interactions, potentially explaining why this modification is not present in native S. cerevisiae U2 snRNA (see Discussion).

**Fig. 2.**
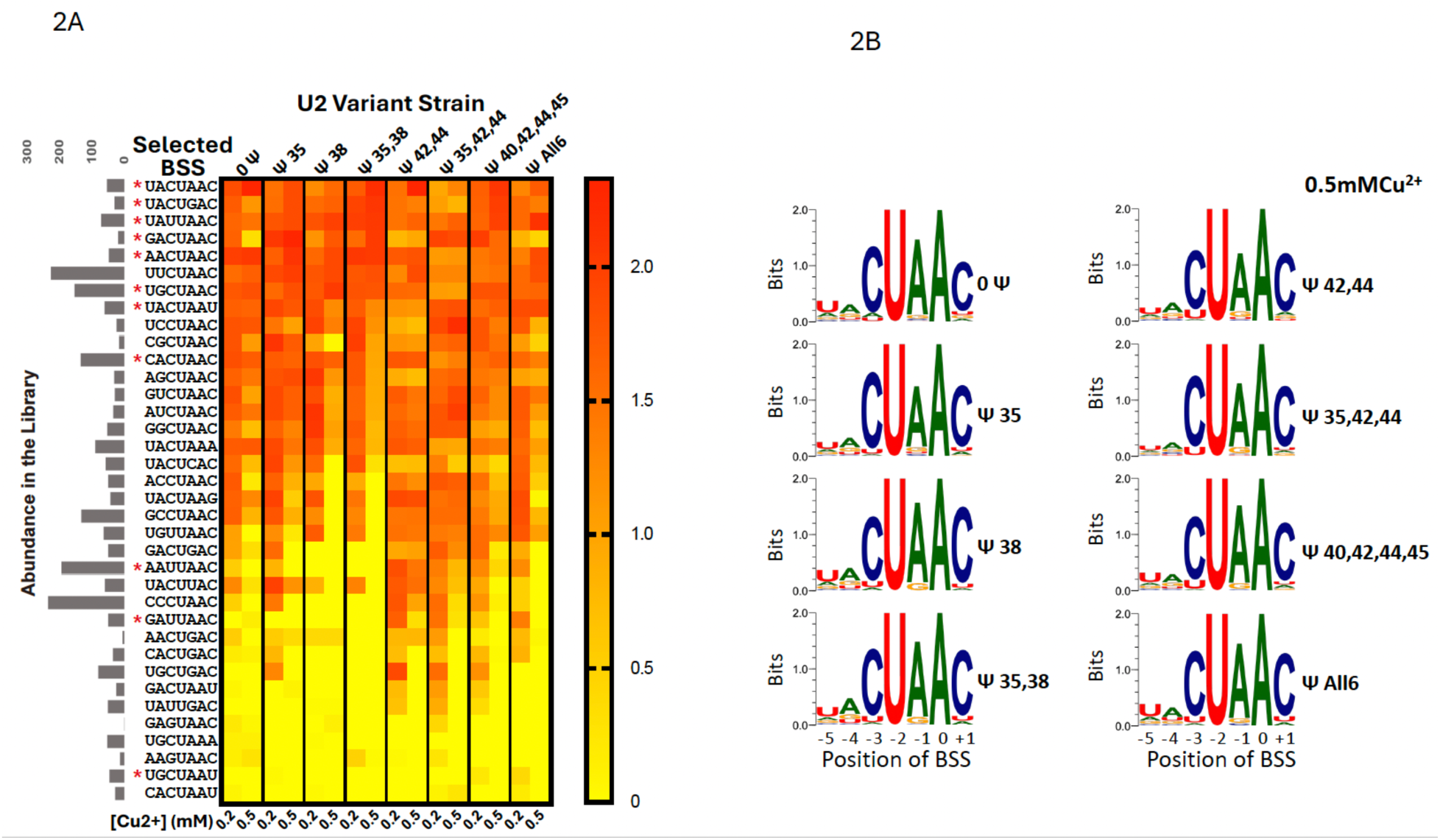
Amplicon sequencing analysis of the BSS library screened across eight U2 Ψ variant strains. (A) U2 BSRR Ψ variant strains (indicated at the top) were individually transformed with the *ACT1-CUP1* reporter library containing randomized BSS sequences. Cells were replica plated onto agar medium containing various concentrations of Cu^2+^. After 3-4 days of incubation, cells grown on plates containing 0.2 mM or 0.5 mM Cu^2+^ (indicated at the bottom) were recovered, plasmids were isolated, and the pre- mRNA BSS region was subjected to amplicon sequencing. All screen-selected BSSs are shown on the left, together with their abundance in the initial library prior to screening. The heatmap depicts the relative abundance of each selected BSS after normalization to the corresponding 0 mM Cu^2+^ control (0.2 mM Cu^2+^/0 mM Cu^2+^; 0.5 mM Cu^2+^/0 mM Cu^2+^). Red asterisks indicate BSSs found in native *S. cerevisiae* genes. Dark red indicates the highest relative abundance, whereas yellow indicates the lowest relative abundance. (B) Consensus sequences of screen-selected BSSs from all eight strains were generated using Shannon entropy analysis.

To validate the screening results (**Fig 2A**), individual *ACT1-CUP1* constructs containing selected BSSs were generated and introduced into each of the eight U2 variant strains. Spot tests were then performed on media containing increasing concentrations of [Cu^2+^] (0 mM, 0.2 mM, 0.5 mM, and 1 mM). As shown in **Supplementary Fig 2**, the spot test results were largely consistent with the screening data (**Fig 2A**). In general, the spot tests recapitulated the finding that a broader range of BSSs was recognized in the Ψ35Ψ42Ψ44, Ψ42Ψ44, and Ψ40Ψ42Ψ44Ψ45 strains, whereas a more limited subset of BSSs was recognized in the Ψ38 and Ψ35Ψ38 strains. Minor discrepancies in BSS recognition were occasionally observed, likely reflecting experimental variation within a very narrow range of splicing efficiencies.

### Ψ in the U2 BSRR contributes to BSS recognition

Upon closer examination of the selected BSSs across the eight U2 variant strains, we found that the recognition efficiency of a given BSS by U2 varied depending on the U2 pseudouridylation pattern, although the magnitude of the effect differed among BSSs. For example, all eleven naturally occurring *S. cerevisiae* BSSs were selected and recognized with varying efficiencies across the different U2 Ψ variant strains (**Fig 2A**). Although the BSS screening results, including those obtained for naturally occurring *S. cerevisiae* BSSs, were confirmed by spot tests (**Supplementary Fig 2**), these assays were performed using the artificial *ACT1-CUP1* reporter system.

To validate these observations in a more physiological context, namely within their native genes, we conducted RT-PCR experiments using RNA recovered from different U2 variant strains to directly measure the splicing efficiencies of naturally occurring pre-mRNAs containing representative BSSs. As shown in **Fig 3A** (left panel) and **Fig 3B**, three genes, each carrying a different BSS, were analyzed in all eight U2 variant strains. Pre-mRNA PTC7, which contains the UGCUAAU BSS, was inefficiently spliced, as indicated by a low mRNA/pre-mRNA ratio (Panel a). This was particularly evident in the 0-Ψ, Ψ38, Ψ35Ψ38, and Ψ35Ψ38Ψ40Ψ42Ψ44Ψ45 (Ψ all 6) strains (lanes 1, 3, 4, and 8, respectively). Pre-mRNA HPC2, carrying the GAUUAAC BSS, exhibited intermediate splicing efficiency (Panel b). Notably, splicing efficiency was lower in the Ψ38 and Ψ35Ψ38 strains (lanes 3 and 4) than in the other U2 variant strains (lanes 1, 2, and 5-8). Pre-mRNA SRB2, which carries the UGCUAAC BSS, was efficiently spliced in all eight U2 variant strains (Panel c). These results are consistent with both the screening data (**Fig 2A**) and spot test results (**Supplementary Fig 2**), in which *ACT1-CUP1* reporters containing the corresponding BSSs exhibited similar U2 variant-dependent differences in recognition (**Fig 3A**, right panel). Taken together, these results demonstrate that distinct U2 BSRR pseudouridylation patterns significantly influence BSS recognition and splicing efficiency.

**Fig. 3.**
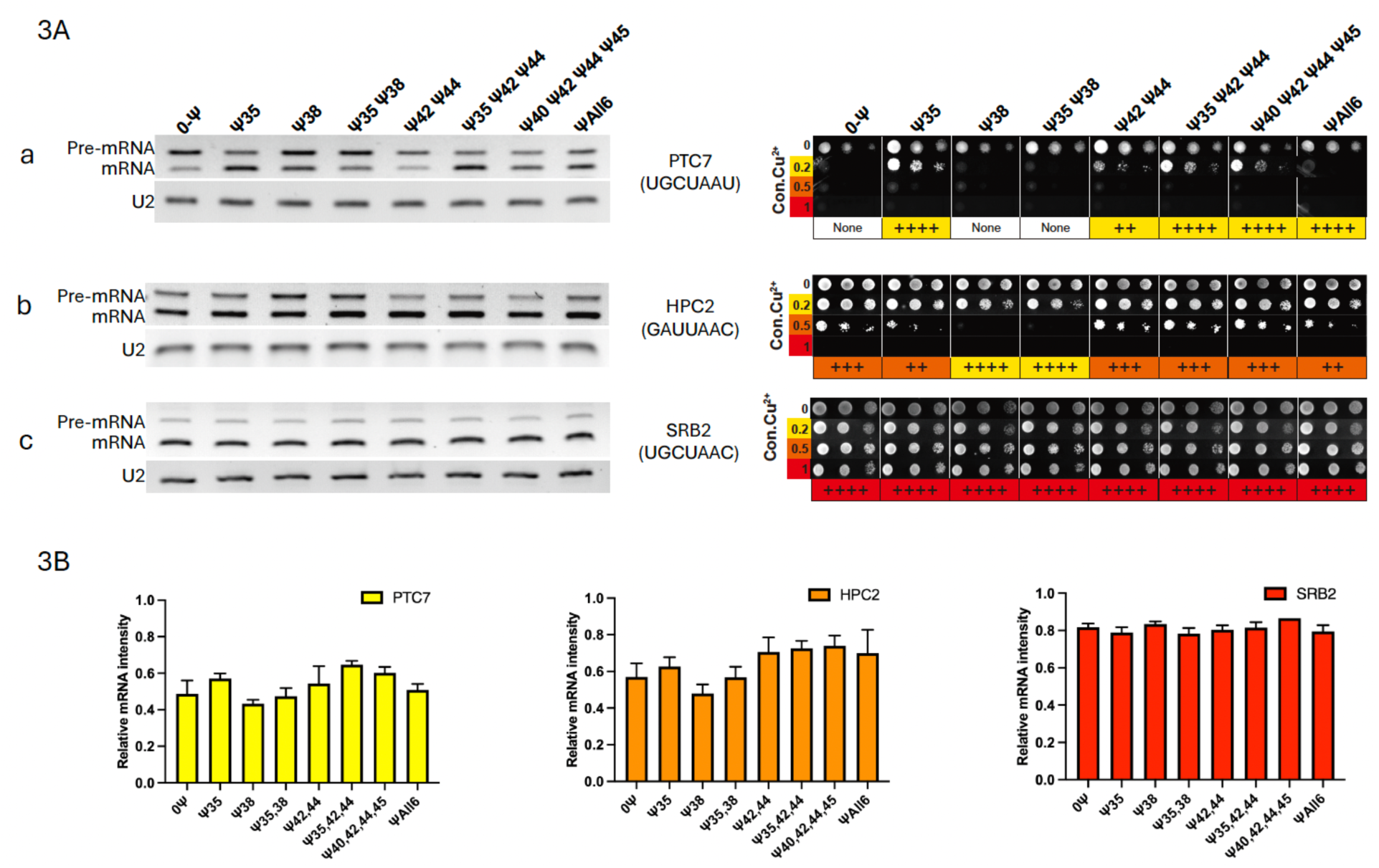
Splicing efficiency of three endogenous yeast pre-mRNAs. (A) Left panel: RT-PCR was used to determine the splicing efficiency of three endogenous pre-mRNAs, PTC7, HPC2, and SRB2, each containing a distinct screen-selected BSS (shown). The three bands correspond to the unspliced transcript (pre-mRNA), spliced transcript (mRNA), and U2 RNA (loading control). Right panel: three *ACT1-CUP1* constructs containing the BSSs corresponding to PTC7, HPC2, and SRB2 were generated. Following transformation of *cup1Δ* U2 Ψ variant strains with the respective *ACT1-CUP1* constructs, cells were plated on media containing the indicated concentrations of Cu^2+^, and splicing efficiency was assessed by growth. Red “+”, highest efficiency; orange “+”, intermediate efficiency; yellow “+”, lowest efficiency; white “none”, no detectable growth. (B) Quantification of the splicing efficiency of endogenous PTC7, HPC2, and SRB2 pre-mRNAs in each strain, based on the RT-PCR data shown in (A). The quantified splicing efficiencies are consistent with the results obtained from the screen.

### Distinct U2 **Ψ** variants can preferentially recognize specific BSSs

From the eight selected BSS lists (**Fig 2A**), we made another interesting observation: for certain BSSs, the order of recognition preference varied substantially among U2 variants and, in some cases, was completely reversed. For example, although several U2 variants, including 0-Ψ, Ψ35, and Ψ35Ψ38, favored UACUAAA over UGUUAAC, the opposite preference was observed in Ψ35Ψ42Ψ44 and Ψ35Ψ38Ψ40Ψ42Ψ44Ψ45.

To confirm these preference orders at the cellular and molecular levels, we transformed the 0-Ψ, Ψ35Ψ42Ψ44, and Ψ35Ψ38Ψ40Ψ42Ψ44Ψ45 (Ψ All 6) strains with the *ACT1-CUP1* reporter gene carrying either the UACUAAA or the UGUUAAC BSS. We then performed a parallel spot test using these transformants. As shown in **Fig 4A**, all strains showed healthy cell growth when transformed with the *ACT1-CUP1* reporter containing the perfect BSS UACUAAC (as a control) (Row 1 of each panel). However, distinct growth patterns were observed when cells were transformed with reporters carrying either UACUAAA or UGUUAAC. Specifically, in the 0-Ψ strain, cells transformed with UACUAAA-*ACT1-CUP1* grew in medium containing 0.5 mM [Cu^2+^] (Panel b), whereas those with UGUUAAC-*ACT1-CUP1* only grew up to 0.2 mM [Cu^2+^]. In contrast, the Ψ35Ψ42Ψ44 and Ψ35Ψ38Ψ40Ψ42Ψ44Ψ45 strains exhibited the opposite pattern: UGUUAAC-*ACT1-CUP1* supported growth at 0.5 mM [Cu^2+^] (Panel a and c), while UACUAAA-*ACT1-CUP1* only supported growth up to 0.2 mM [Cu^2+^].

**Fig. 4.**
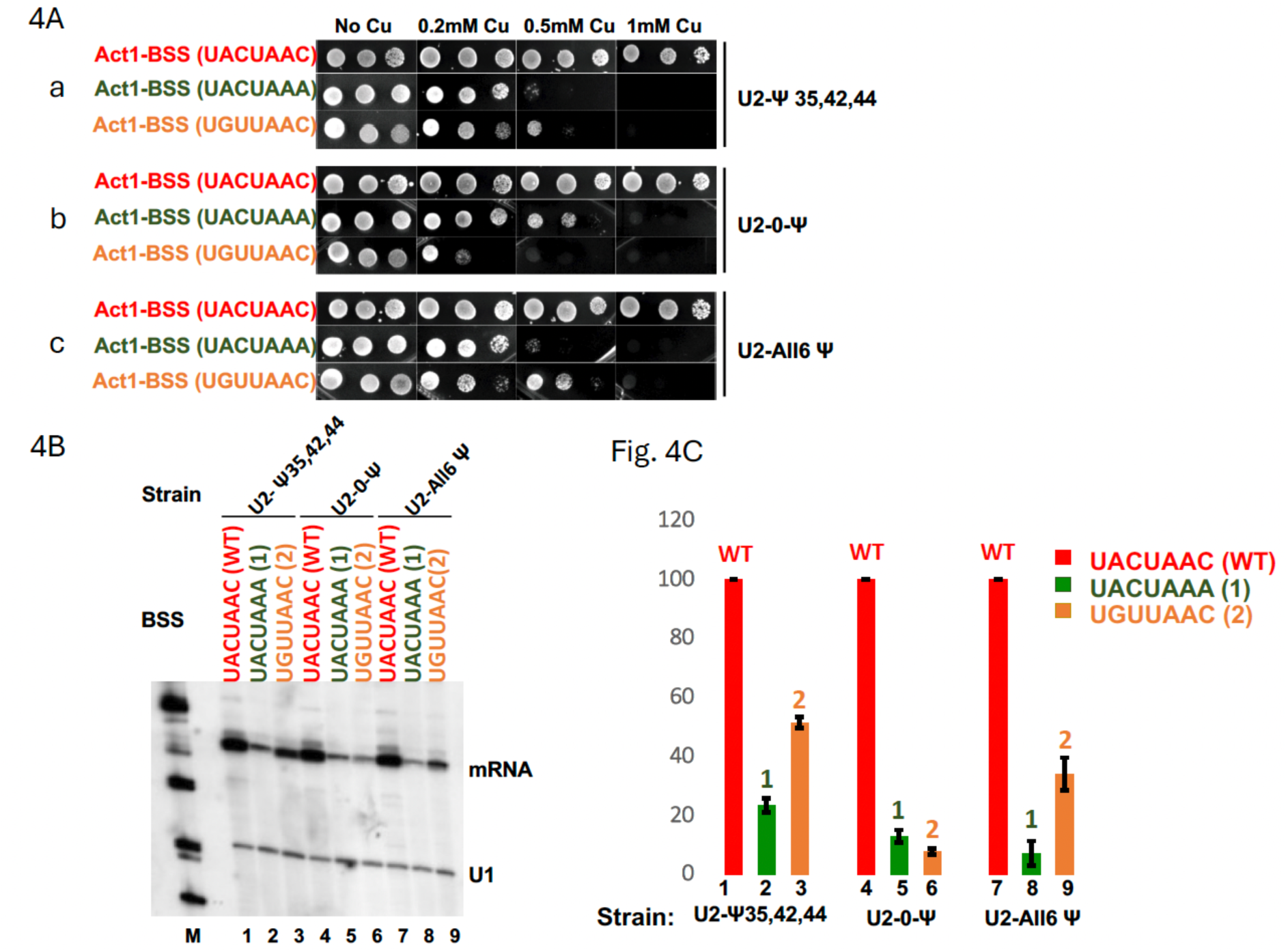
Validation of differential splicing of pre-mRNAs containing distinct BSSs in U2 Ψ variant strains. (A) *ACT1-CUP1* reporters containing either a UACUAAC, UACUAAA, or UGUUAAC BSS were constructed. After transformation of the indicated U2 Ψ variant strains with each reporter, spot assays were performed on media containing indicated concentrations of Cu^2+^. (B) Total RNA was isolated from the cells shown in (A), and primer-extension analysis was performed to assess splicing efficiency. Bands corresponding to the spliced mRNA and U1 RNA (loading control) are indicated. (C) Quantification of the splicing efficiency of each *ACT1-CUP1* reporter based on the data shown in (B). Lane numbers correspond to those shown in (B). Data represent three biological replicates (n=3). Statistical significance was determined using a two-sided unpaired Student’s *t*-test.

In parallel, we extracted total RNA from these transformed strains and performed primer-extension analysis to measure splicing efficiency directly. As shown in **Fig 4B and 4C**, efficient splicing was observed in all strains transformed with the perfect BSS (UACUAAC)-containing *ACT1-CUP1* gene (lanes 1, 4, and 7). In contrast, splicing of pre-mRNAs containing either the UACUAAA or UGUUAAC BSS was less efficient. More specifically, in 0-Ψ cells, UACUAAA pre-mRNA was spliced slightly more efficiently than UGUUAAC pre-mRNA (compare lane 5 with lane 6). In contrast, in the Ψ35Ψ42Ψ44 and Ψ35Ψ38Ψ40Ψ42Ψ44Ψ45 strains, UGUUAAC pre-mRNA was spliced more efficiently than UACUAAA pre-mRNA (compare lane 2 with lane 3, and lane 8 with lane 9). These results, together with the screening data (**Fig 2A**) and spot test results (**Fig 4A**), demonstrate that distinct U2 pseudouridylation patterns can alter, and in some cases completely reverse, BSS recognition preferences.

### Distinct U2 Ψ variants differentially select competing BSSs and influence alternative splicing

The differential preference described above led us to predict that, when two BSSs (e.g., UACUAAA and UGUUAAC) are placed side by side in the same pre-mRNA, different U2 variant strains would use them differently. Specifically, strains such as 0-Ψ, Ψ35, and Ψ35Ψ38 were predicted to preferentially use UACUAAA, whereas the Ψ35Ψ42Ψ44, Ψ40Ψ42Ψ44Ψ45, and Ψ35Ψ38Ψ40Ψ42Ψ44Ψ45 strains were predicted to favor UGUUAAC.

To test this hypothesis, we constructed two versions of the *ACT1-CUP1* reporter, containing both BSSs in tandem. In Construct 1 (C1), UACUAAA precedes UGUUAAC; in Construct 2 (C2), the order is reversed (UGUUAAC followed by UACUAAA) **(Fig 5A)**. Each construct was transformed into four U2 variant strains (0-Ψ, Ψ35Ψ42Ψ44, Ψ40Ψ42Ψ44Ψ45, and Ψ35Ψ38Ψ40Ψ42Ψ44Ψ45). We then extracted RNA and performed primer-extension, which stops at the branch point of the lariat structure, to determine which BSS was utilized during splicing. As shown in **Figs 5B and 5C**, the 0-Ψ strain consistently preferred UACUAAA, regardless of which construct (C1 or C2) was used (lanes 3 and 4). In contrast, the Ψ35Ψ42Ψ44, Ψ40Ψ42Ψ44Ψ45, and Ψ35Ψ38Ψ40Ψ42Ψ44Ψ45 strains all favored UGUUAAC, again independent of construct order (lanes 1, 2, 5-8).

**Fig. 5.**
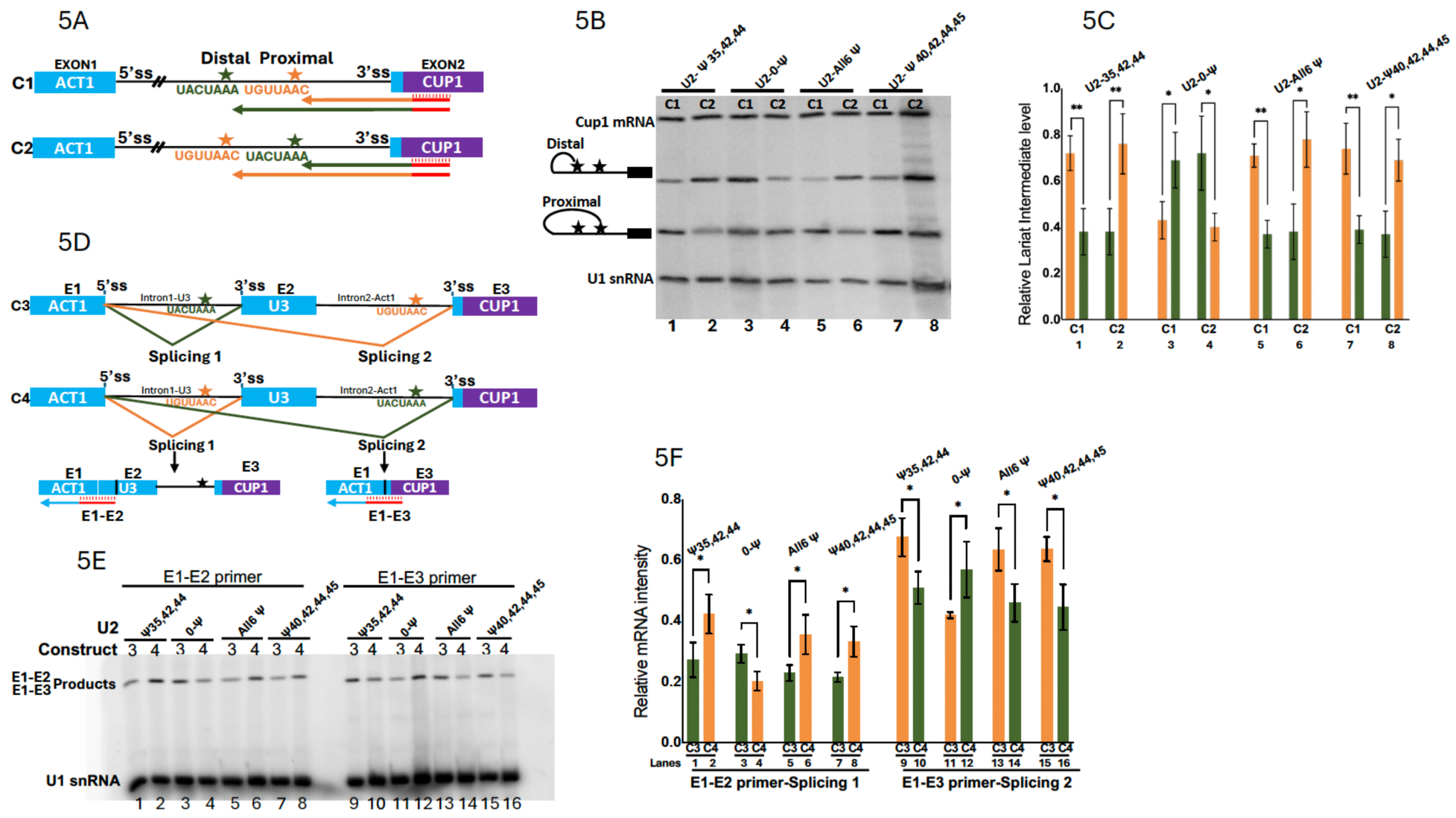
Differential recognition of BSSs by distinct U2 Ψ variants during splicing. (A) Two *ACT1-CUP1* constructs containing tandem BSSs were generated: C1, with UACUAAA (green) upstream of UGUUAAC (brown), and C2, with UGUUAAC upstream of UACUAAA. Stars indicate the branchpoint nucleotides, which form the lariat intermediate after the first step of splicing and cause primer extension termination. The primer used for analysis is indicated by the red lines. (B) Primer-extension analysis detected preferential usage of either UACUAAA or UGUUAAC branch site (indicated) during splicing in the indicated U2 Ψ variant strains. The U1 primer-extension product is shown as a loading control. (C) Quantification of branch-site usage based on the data shown in (B). Lane numbers correspond to those shown in (B) Data represent three biological replicates (n=3). Statistical significance was assessed using a two-sided unpaired Student’s *t*-test. \**P* < 0.05, and \*\**P* < 0.01. (D) Two additional constructs (C3 and C4) containing tandem BSSs (UACUAAA in green and UGUUAAC in brown), each associated with its own 3’ splice site, were generated. The possible alternative splicing pathways are illustrated. E1-E2 and E1-E3 exon-exon junction primers were used in primer-extension assays to quantify the relative levels of alternatively spliced products. (E) Total RNA was isolated from the indicated U2 Ψ variant strains, and primer-extension analyses were performed using the E1-E2, E1-E3, and U1 primers. The corresponding extension products are indicated. (F) Relative levels of the E1-E2 and E1-E3 primer-extension products were quantified after normalization to the U1 primer-extension product. Lane numbers correspond to those shown in (E). Data represent three biological replicates (n=3). Statistical significance was determined using a two-sided unpaired Student’s *t*-test. \**P* < 0.05.

To further analyze whether the U2 BSRR Ψs directly influence BSS choice in an alternative-splicing-like context, we modeled our constructs after the *Drosophila Transformer* pre-mRNA (Sosnowski et al. 1989; Tian and Maniatis 1993; Valcarcel et al. 1993; Black 2003) and created two additional constructs (C3 and C4), each containing two distinct BSSs linked to their own (and identical) 3’ splice sites (**Fig 5D).** Constructs 3 and 4 were conceptually similar to the *Drosophila Transformer* system, but differed in one important respect. In *Transformer*, competing 3’ splice sites are distinguished primarily by differences in their upstream polypyrimidine tracts (Valcarcel et al. 1993). In contrast, the competing splice sites in Constructs 3 and 4 differed only in their BSS sequences. Specifically, Construct 3 contained a UACUAAA BSS linked to a 3’ splice site followed by a UGUUAAC BSS linked to an identical 3’ splice site. Construct 4 contained the same elements in the opposite order.

To detect the splicing products, we performed primer-extension initially using several primers complementary to the common 3’ exon. However, these experiments yielded several unexplained high-background bands, complicating interpretation. To overcome this issue, we switched to two exon-exon junction primers. Although the use of two independent exon-exon junction primers prevented direct comparison of absolute alternative splicing efficiencies, differences in the relative levels of primer-extension products generated from Constructs 3 and 4 nevertheless provided a reliable measure of BSS preference in different U2 variant strains. Specifically, the 0-Ψ strain consistently produced a stronger signal for UACUAAA usage than for UGUUAAC usage, regardless of the construct (C3 or C4) and primer (Exon 1-Exon2 or Exon 1-Exon 3) employed (**Figs 5D-F**, lanes 3, 4, 11, and 12; and compare the ratio of pair lanes 3 and 4 with the ratio of pair lanes 11 and 12). In contrast, stronger signals corresponding to UGUUAAC usage than to UACUAAA usage were detected in the Ψ35Ψ42Ψ44, Ψ40Ψ42Ψ44Ψ45, and Ψ35Ψ38Ψ40Ψ42Ψ44Ψ45 strains (**Figs 5D-F**, lanes 1, 2, 5-10, and 13-14; and compare the ratio of each pair). Collectively, these experiments demonstrate that distinct U2 BSRR pseudouridylation patterns can alter BSS choice. Whereas the 0-Ψ U2 variant preferentially utilized UACUAAA, U2 variants containing Ψ35Ψ42Ψ44, Ψ40Ψ42Ψ44Ψ45, or Ψ35Ψ38Ψ40Ψ42Ψ44Ψ45 preferentially utilized UGUUAAC. These differences in BSS selection directly influenced alternative splice-site usage.

### Ψ in the U2 BSRR influences the recruitment of Prp5 to modulate BSS recognition

To understand how U2 Ψs influence BSS selection, we focused on Prp5, an ATPase that recognizes and monitors the U2-BSS duplex during early spliceosome assembly (Xu and Query 2007) and preferentially associates with U2 pseudouridylated at specific positions within the branchpoint-interacting stem loop (BSL) (Liang and Cheng 2015; Wu et al. 2016). We employed a plasmid-shuffling strain in which chromosomal *PRP5* was replaced by a plasmid-borne wild-type *PRP5* marked with *URA,* thus allowing selection against the plasmid under 5-FOA (Xu and Query 2007).

Synthetic functional interaction analysis was performed across all eight U2 variant strains (**Fig 6A**, lanes 1-8). Replacing the wild-type *PRP5* plasmid with either an identical wild-type PRP5 plasmid (Row b) or a fully functional Prp5 mutant carrying a single N399D substitution (Row f) had no detectable effect on growth in all U2 variant strains. In contrast, replacing the wild-type *PRP5* plasmid with a partially functional prp5 mutant, in which the conserved motif III SAT tripeptide was mutated to GAR, resulted in severe growth defects in the 0-Ψ, Ψ35, Ψ38, and Ψ35Ψ38 strains (Row d, lanes 1-4), a slight growth defect in the Ψ35Ψ38Ψ40Ψ42Ψ44Ψ45 strain (Row d, lane 8), and almost no defects in the Ψ42Ψ44, the Ψ35Ψ42Ψ44, and the Ψ40Ψ42Ψ44Ψ45 strains (lanes 5-7). These results are consistent with our findings discussed above, which show that distinct U2 BSRR pseudouridylation patterns differentially influence BSS recognition, and further suggest that Ψ42 and Ψ44 enhance Prp5-mediated monitoring of U2-BSS interactions by promoting Prp5 recruitment (Wu et al. 2016). These results are also consistent with our findings that Ψ38 generally played an inhibitory role in BSS recognition and pre-mRNA splicing (**Fig 2**), possibly by inhibiting Prp5 recruitment.

**Fig. 6.**
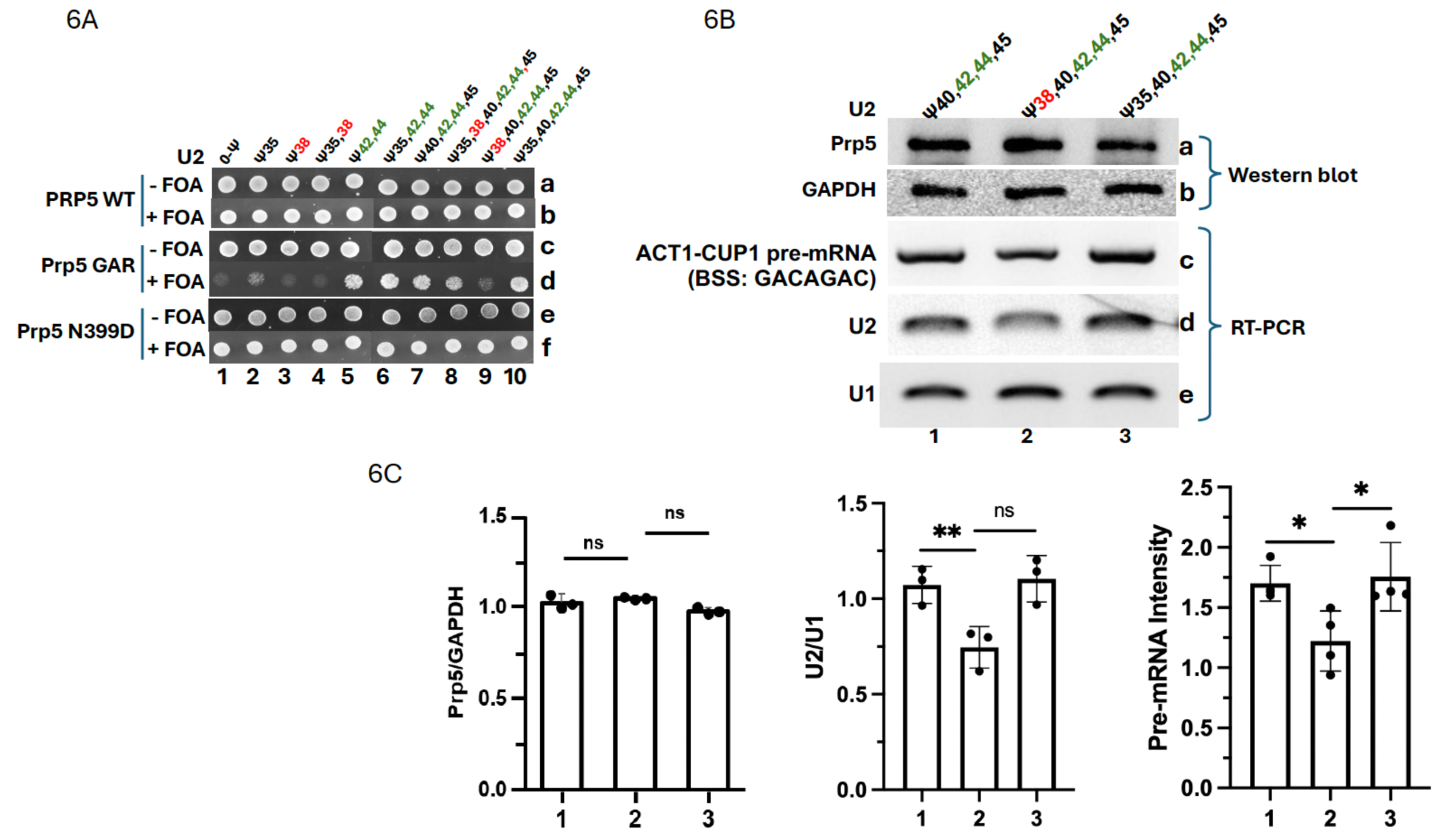
Synthetic lethality analysis of U2 Ψ variants and Prp5 mutants, and characterization of Prp5-U2 and Prp5-U2-pre-mRNA interactions. (A) Cells containing a plasmid-borne *PRP5* gene marked with URA3 were transformed with plasmids expressing either wild-type Prp5 (rows a and b), Prp5-GAR (rows c and d), or Prp5 N399D (rows e and f) (Xu and Query 2007), together with plasmids encoding designer gRNAs targeting U2 at the indicated positions shown at the top. Transformants were grown on medium lacking 5-fluoroorotic acid (-5FOA) or containing 5-fluoroorotic acid (+5FOA), as indicated. When combined with the wild-type Prp5 (row b) or Prp5 N399D (row f), all U2 Ψ variant strains showed no growth defects (lanes 1-10). In contrast, when combined with Prp5-GAR, only the U2 Ψ variant strains containing Ψ42 and Ψ44 grew well (row d, lanes 5-8). For example, the U2 Ψ40Ψ42Ψ44Ψ45 strain (lane 7) showed strong growth. However, addition of Ψ38 (lane 9), but not Ψ35 (lane 10), to the parental U2 Ψ40Ψ42Ψ44Ψ45 strain resulted in loss of viability (compare lanes 9 and 10 with lane 7). (B) Prp5-U2 and Prp5-U2-pre-mRNA interactions were analyzed in the Ψ40Ψ42Ψ44Ψ45, Ψ38Ψ40Ψ42Ψ44Ψ45, and Ψ35Ψ40Ψ42Ψ44Ψ45 strains. Cells expressing FLAG-tagged wild-type Prp5 were co-transformed with gRNA plasmids directing U2 pseudouridylation to generate the Ψ40Ψ42Ψ44Ψ45, Ψ38Ψ40Ψ42Ψ44Ψ45, or Ψ35Ψ40Ψ42Ψ44Ψ45 variants, and an *ACT1-CUP1* reporter plasmid containing the weak BSS GACAGAC. Cell lysates were subjected to anti-FLAG immunoprecipitation (Prp5 is FLAG-tagged). Precipitated samples were analyzed by anti-FLAG western blotting (row a). As a control, cell lysates were also analyzed by anti-GAPDH western blotting (row b). In parallel, *ACT1-CUP1* pre-mRNA (row c), U2 (row d), and U1 (row e) co-immunoprecipitated with FLAG-tagged Prp5 were analyzed by RT-PCR. (C) Quantification of co-immunoprecipitated Prp5 (normalizing to GAPDH), ACT1-CUP1 pre- mRNA (normalizing to U1), and U2 snRNA (normalizing to U1) based on the data shown in (B). Lane numbers correspond to those shown in (B). Data represent three or four biological replicates (n ≥ 3). Statistical significance was determined using a two-sided unpaired Student’s *t*-test. \**P* < 0.05; \*\**P* < 0.01.

To test whether Ψ38 inhibits Prp5 recruitment and further investigate the molecular mechanism underlying BSRR Ψ function, we generated two additional strains by adding Ψ38 or Ψ35 to the Ψ40Ψ42Ψ44Ψ45 variant. Remarkably, although the new Ψ38Ψ40Ψ42Ψ44Ψ45 strain showed no defect when combined with either wild-type or N399D Prp5 (**Fig 6A**, Rows b and f, lane 9), it exhibited a severe synthetic growth defect when combined with the Prp5 GAR mutant (**Fig 6A**, Row d, lane 9), an observation that contrasts sharply with that of the parental Ψ40Ψ42Ψ44Ψ45 strain (lane 7; compare lane 7 with lane 9). In contrast, the other new strain, Ψ35Ψ40Ψ42Ψ44Ψ45, exhibited no growth defect and even showed slightly improved growth compared with the parental Ψ40Ψ42Ψ44Ψ45 strain when combined with the Prp5 GAR mutant (Row d, compare lane 10 with lane 7).

To directly examine whether Ψ38 negatively impacts Prp5 recruitment, we performed Prp5-U2 co-immunoprecipitation analysis (**Figs 6B and 6C**). Anti-FLAG antibody efficiently precipitated similar amounts of FLAG-tagged Prp5 from all three strains (Row a, lanes 1-3), suggesting comparable Prp5 expression levels. Anti-FLAG antibody also co-precipitated the parental Ψ40Ψ42Ψ44Ψ45 U2 variant (Row d, lane 1), whereas inclusion of Ψ38 significantly reduced U2 recovery (Row d, lane 2). By contrast, addition of Ψ35 had no effect (and perhaps a slightly stimulating effect) on Prp5 binding and co-precipitation (Row d, lane 3), in agreement with the synthetic functional interaction results (Fig 6A). No differences were detected for U1 controls (Row e, lanes 1-3). Consistent results were obtained from Prp5-U2-pre-mRNA co-precipitation assays using an inefficiently spliced pre-mRNA substrate with a very weak BSS (GACAGAC): pre-mRNA recovery was reduced with the Ψ38-containing U2 variant (Ψ38Ψ40Ψ42Ψ44Ψ45) (Row c, lane 2), but high and comparable between Ψ35-containing (Ψ35Ψ40Ψ42Ψ44Ψ45) (Row c, lane 3) and parental control (Ψ40Ψ42Ψ44Ψ45) (Row c, lane 1) variants. These results indicate that pseudouridylation at position 38 has an inhibitory effect on Prp5 binding, opposite to Ψ42 and Ψ44, which promote Prp5 binding (Wu et al. 2016). The close correspondence between the binding results (Figs 6B and 6C) and the genetic functional phenotypes (Fig 6A) suggests that Ψ38, together with Ψ42, Ψ44, and other BSRR Ψs, modulates BSS recognition by differentially regulating Prp5 recruitment to the U2 and U2-pre-mRNA complexes.

## Discussion

By conducting screening experiments in which pre-mRNAs with randomized BSSs were spliced in various U2 Ψ variant strains, we identified sets of BSSs that were differentially recognized by specific U2 variants. These results indicate that Ψs within the U2 BSRR differentially contribute to BSS selection. We measured endogenous BSS recognition and obtained results that were consistent with the screening data. By performing synthetic lethality assays using mutant Prp5 and U2 Ψ variants, together with Prp5-U2-pre-mRNA binding assays, we further showed that U2 Ψs, especially Ψ38, Ψ42, and Ψ44, play important roles in Prp5 recruitment and BSS recognition.

The differential recognition of different BSSs by distinct U2 Ψ variants suggests that individual Ψs can influence branch-site choice and may contribute to alternative splicing. To test this hypothesis, we created pre-mRNA constructs containing two BSSs, either linked to a common 3’ splice site or each linked to its own 3’ splice site. Distinct splicing patterns emerged depending on the U2 Ψ variant, reinforcing the idea that U2 Ψs possess the capacity to influence alternative splicing through differential branch-site selection when alternative branch-site choices are available. Although our experiments were performed in yeast, the effects of Ψs are likely to be similar in higher eukaryotes, given the conserved mechanism of pre-mRNA splicing (Kastner et al. 2019; Plaschka et al. 2019); however, additional regulatory elements, including splicing factors, likely further modulate alternative splicing outcomes, especially in higher eukaryotes. The combined action of these components ultimately determines splicing patterns.

Previously, we demonstrated that Ψ42 and Ψ44 of the 3’ half of the U2 BSRR are critical structural elements recognized by Prp5 (Wu et al. 2016). In the current study, we extended these findings by showing that other Ψs in the U2 BSRR, including at least Ψ38 in the 5’ half of the BSRR, also act as recognition elements. Remarkably, we found that Ψ38 reduces Prp5 binding, whereas Ψ42 and Ψ44 promote Prp5 binding (**Fig 7**). Consistent with this model, analyses combining U2 Ψ variation with the Prp5 mutation revealed a strong correlation between BSS recognition efficiency and the functional and physical interactions among pre-mRNA, U2 Ψ variants, and Prp5. These findings support a model that U2 BSRR Ψs constitute important molecular determinants of the U2-BSS duplex recognized by Prp5 during spliceosome assembly (**Fig 7**).

**Fig. 7.**
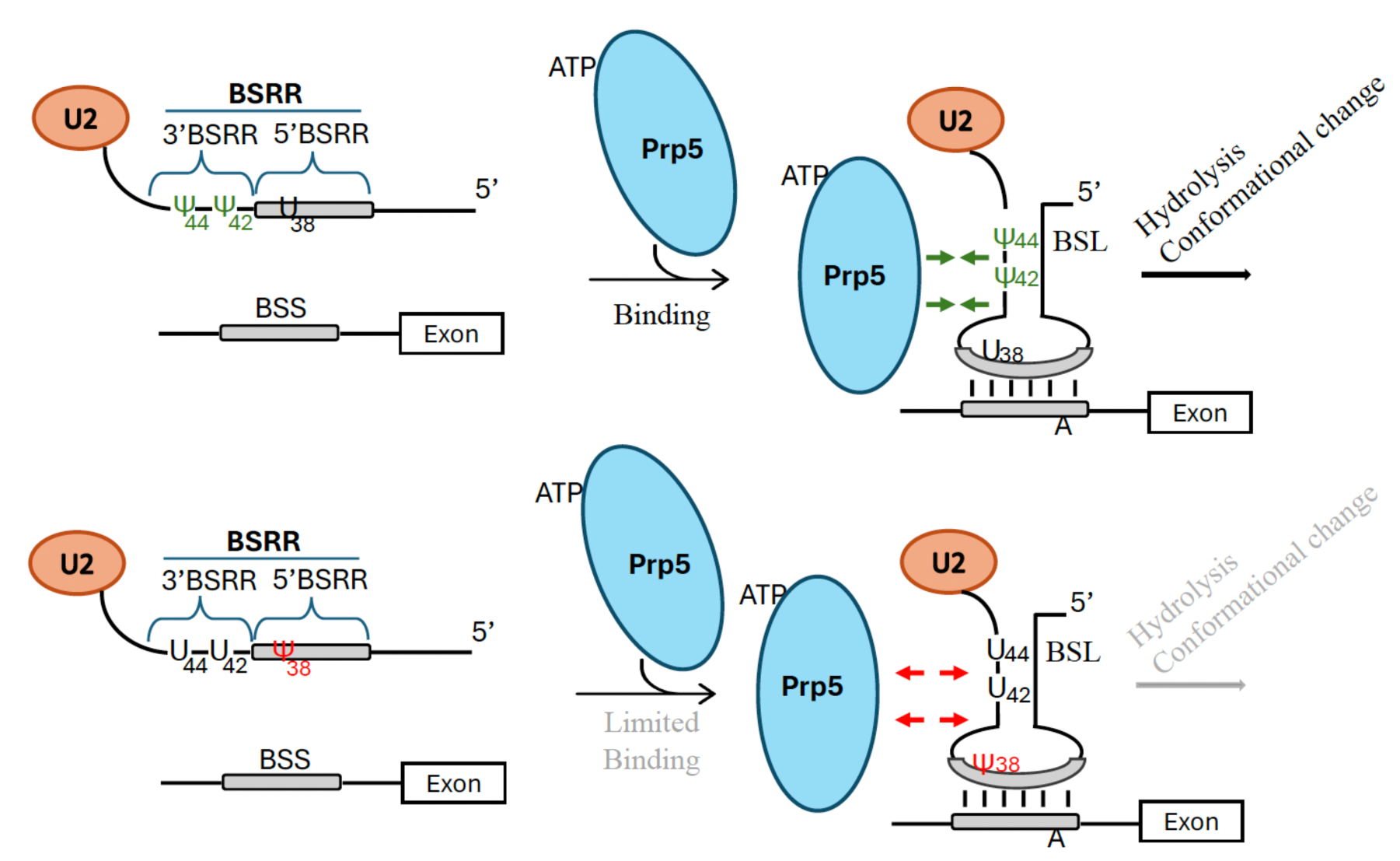
Model illustrating the contributions of pseudouridines in the U2 BSRR to modulation of Prp5 binding and recruitment. Prp5 is known to bind to U2 snRNA through the BSL structure (Perriman and Ares 2010; Liang and Cheng 2015), thereby stimulating its ATPase activity and promoting a conformational rearrangements that facilitate Complex A formation. In the U2 U38Ψ42Ψ44 background, Ψ42 and Ψ44 enhance recruitment of Prp5 to the U2 BSL, promoting Prp5 activity and subsequent Complex A formation. In contrast, in the U2 Ψ38U42U44 background, Ψ38 inhibits Prp5 binding, either directly or indirectly, resulting in reduced Prp5 activity and less efficient Complex A formation.

How might Ψ influence Prp5 recruitment to U2 and the recognition of the U2-BSS duplex by Prp5? One possibility is that Ψ alters the local structure of the U2-BSS duplex, thereby influencing its recognition by Prp5 (Arnez and Steitz 1994; Charette and Gray 2000; Newby and Greenbaum 2002; Wu et al. 2016). Alternatively, Ψ could directly promote or inhibit Prp5 binding. The additional hydrogen-bond donor provided by Ψ, together with differences in base rigidity relative to uridine (Charette and Gray 2000; Karijolich and Yu 2010), may facilitate or impede specific contacts with Prp5. To further investigate these possibilities, high-resolution structural analyses, such as cryo-EM, will be essential to reveal the precise molecular interactions involved.

In light of these findings, it is notable that a single U2 snRNA sequence contains multiple Ψs, yet exhibits only 80-90% pseudouridylation at any given Ψ site (Zhao and Yu 2004a; Borchardt et al. 2020). This partial modification suggests the presence of multiple U2 sequence variants within a cell. Such diversity may be functionally significant, particularly in higher eukaryotes, where the greater diversity of branch-site sequences may require a broader repertoire of U2 snRNA variants. Importantly, Ψ exhibits greater pairing versatility than uridine (U), including an enhanced capacity to stabilize noncanonical base-pairing interactions (Kierzek et al. 2014; Pan et al. 2026), a property that may increase the adaptability of U2 snRNA in recognizing diverse BSSs. This increased pairing flexibility, combined with sequence variability, likely contributes to the complexity and precision of splicing regulation, especially in higher eukaryotes with more intricate transcriptomes. The fact that some Ψs (such as Ψ38) negatively impact BSS recognition suggests an additional layer of complexity in splicing regulation, in which certain pre-mRNAs may require highly efficient splicing, while others may not, depending on cellular needs. In contrast, S. cerevisiae, which encounters far fewer BSS variants (more than 90% conform to the optimal sequence UACUAAC), maintains relatively fewer U2 Ψ variants (only three Ψs). Given its highly conserved BSS and the absence of Ψ38 from the yeast U2 BSRR, these observations suggest that S. cerevisiae has evolved to favor highly efficient branch-site recognition and splicing.

## Materials and Methods

### Construction of the Act1-Cup1 intron libraries

The *ACT1-CUP1* reporter plasmid was used to build up the BSS-randomization library. The 463 bp, double-stranded library sequence containing partial *ACT1* exon, intron, and full length of *CUP1* coding sequence (Lesser and Guthrie 1993; Wu et al. 2016) was synthesized from Invitrogen^TM^ GeneArt^TM^ Strings^TM^ DNA Libraries that the 6 BSS nucleotides (except for the branchpoint nucleotide adenosine) were randomly mutated (25% A, 25% T, 25% C, 25% G), with XhoI and BglII restriction sites at the 5’ end and 3’ end, respectively. The Restriction enzyme-digested fragments were inserted into the *ACT1-CUP1* reporter plasmid (YEplac195, URA3, 2μ, GPD promoter). The ligation products were transformed into *E.coli* (XL-10 Gold) and multiplied to prepare the *AC1-CUP1* plasmid libraries (**Fig 1**).

### Construction of the guide RNA targeting U2 pseudouridines and yeast transformation

10 U2 BSRR variant strains were created, each with a distinct pseudouridylation pattern within the BSRR. All of these strains were created from the 0 Ψ strain named 6 GB (*MATa cup1::ura3, pus7::HIS, snR81::KAN, pus1::TRP, leu2 Δ0, ura3 Δ0, trp1 Δ0, lys2 Δ0, ade2 Δ0, GAL+ his3 Δ0*) (Wu et al. 2016), whose parental strain is YCL46 (Lesser and Guthrie 1993). Cells were transformed with a plasmid carrying different PugU2 box H/ACA RNA sequences (YEplac181, LEU2, 2μ, GPD promoter) (**Supplementary Table 1**) (Wu et al. 2016), where the guide pocket sequences were altered to target the corresponding uridine site in the U2 BSRR. In the strains with a single Ψ, such as Ψ35, one box H/ACA RNA with both pocket sequences targeting the same site U35, while in the strains with two Ψs, such as Ψ42, 44, one box H/ACA RNA with two pocket sequences targeting U42 and U44, respectively. For a BSRR with more than two Ψs, two or three box H/ACA RNA sequences were inserted into the corresponding restriction sites (**Supplement Table 1**). The constructed plasmids were transformed into *E. coli* (XL-10 Gold) and amplified. After being purified from *E. coli*, the plasmids were subsequently transformed into 0 Ψ yeast cells (6GB), and positive colonies were selected on synthetic-defined (SD) solid media lacking Leucine. Further confirmation of the correct pseudouridines was done by pseudouridylation assay [CMC-primer extension analysis (see below)] using total RNA isolated from transformed yeast cells (see below).

### RNA isolation

Yeast cells were cultured in 10 ml of synthetic defined (SD) medium. The cells were harvested around OD 1.0-1.5 and spun at 4700 rpm for 10 min. The cell pellets were vigorously vortexed with 200 μl of 0.5 mm acid-washed glass beads (BioSpec product, Cat.11079105) and 1ml TRIzol (Invitrogen) in a bead beater (30s, four times on ice). 200 μl of chloroform was added to the same 2 ml screw tube, and the tubes were vigorously inverted for 1 min. The lysates were centrifuged at 13,000 rpm for 10 min, and the upper aqueous layer was extracted twice with 200 μl of PCA (Phenol/chloroform/isoamyl alcohol [25:24:1]). RNA in the aqueous phase was precipitated with 100% ethanol. After washing the RNA pellet with 75% ethanol, the pellet was air-dried and dissolved in an appropriate volume of dd H_2_O.

### Pseudouridylation assay

After the extraction of total RNA, CMC-modification followed by primer-extension was conducted. 8-12 μg of total cellular RNA extracted above was dissolved in 20 μl of ddH_2_O. In the CMC minus group, 80 μl BEU buffer (7 M Urea, 4 mM EDTA, pH 8.0∼8.5, 50 mM Bicine) without CMC, plus 20 μl of ddH_2_O, was added, whereas in the CMC plus group, 80 μl of BEU buffer, plus 20 μl of 1 M CMC solution, was added (Final 0.2 M CMC solution in BEU buffer). The mixture was incubated at 37°C for 20 min. RNA was ethanol-precipitated and washed with 70% ethanol. The CMC-treated RNA pellet was resuspended in 50 μl of 100 mM sodium carbonate, pH 10.4, and incubated at 37℃ for 2 hours. RNA was extracted by PCA, ethanol-precipitated, and washed with 70% ethanol. The precipitated RNA pellet was used for primer extension analysis. Briefly, the CMC-treated RNA pellet was resuspended in 4.5 μl of ddH_2_O and mixed with 1.6 μl of 5x annealing buffer (250 mM Tris-HCl, pH 8.3, 300 mM NaCl, 50 mM DTT) and 1 μl (100 nM) of 5’ ^32^P-labeled primer 5’TTGAGGTCATTTCAGTTGTTAC3’, complementary to nucleotides 55-76 of U2 snRNA (for mapping Ψs within the U2 BSRR). The mixture was heated at 95 °C for 3 min and immediately placed at 55 ℃ for 30 min to allow the annealing of primer to RNA. The mixture was then cooled down on ice, and 12 μl of reverse transcription (RT) mixture (18 mM Tris-HCl, pH 8.3, 21 mM NaCl, 3.6 mM DTT, 11 mM Mg(OAc)_2_, 0.72mM each dNTP, and 0.25 units of AMV reverse transcriptase (Promega)) was added. The reverse transcription reaction was carried out at 37℃ for 5 min followed by another 30 min at 42 ℃. The reaction was stopped by adding G50 buffer (20 mM Tris-HCl, pH 7.5, 300 mM sodium acetate, 2 mM EDTA, and 0.2% sodium dodecyl sulfate) and ethanol precipitation. The pellet was resuspended in ddH_2_O and analyzed by 8% denaturing gel electrophoresis and autoradiography.

### Library plasmids transformation and copper screening

Yeast cells with different Ψ (SD Leu) were transformed with library plasmids, and colonies were selected on SD Leu/Ura solid media. Briefly, for each strain, 800 ng of plasmid DNA was transformed into 3 ml of early log-phase cells (OD between 0.5 and 1.0). Multiple plates were prepared to obtain enough colonies. After being mixed with 300 μl of OST buffer (240 μl of 40% (w/v) PEG3350, 36 μl of 1M lithium acetate, 36 μl of 1M DTT), the transformants were incubated at 42°C for 40 min. Cells were then spun down at 5000 rpm for 3 min. The OST buffer was removed completely, and the cell pellets were resuspended in 200 μl of SD Leu/Ura liquid media and spread on SD Leu/Ura selective agar plates. Cells were then replica-plated onto copper-containing agar plates at [Cu^2+^] concentrations of 1 mM, 0.5 mM, 0.2 mM, and 0 mM. The positive colonies appeared after 3-4 days at 30°C. The copper-screened colonies were harvested and rinsed with ddH_2_O. Library plasmids were extracted with Zymoprep Yeast Plasmid Miniprep II kit (D2004). Amplicons for sequencing were prepared by PCR. For every PCR reaction, a total volume of 25 μl was applied, including Forward primer (1.25 μl) 5’**ACACTCTTTCCCTACACGACGCT-CTTCCGATCT**GACTGATCTGTAATAACCACG3’ (Bold letters are the Illumina Adapter Sequences), Reverse primer (1.25 μl) 5’**GACTGGAGTT-CAGACGTGTGCTCTTCCGATCT**GGCATTGGCACTCATGACCTTC3’, Q5® High-Fidelity 2X Master Mix (M0492S, 12.5 μl), and Template 200 ng (10 μl). After gel purification and extraction, the PCR products (307 bp) from different sample groups were brought to a concentration of 100 ng/μl with ddH_2_O, and 20 μl of the sample was sent for sequencing.

### Spot test assay

To validate the screening results, all *ACT1-CUP1* reporter genes with screen-selected BSSs were constructed and separately transformed into the *cup1-deletion* strain. Each yeast isolate was grown in SD Leu/Ura selective media to an OD of 1.0. Cultures were then serially diluted (10×) twice with ddH_2_O. 5 μl from each dilution was spotted on SD Leu/Ura selective media plates with varying copper concentrations. Plates were then incubated at 30°C for 2-4 days and photographed.

### RT-PCR for measuring the splicing efficiency of endogenous pre-mRNAs

Total RNA was extracted from eight strains with different U2 BSRR pseudouridylations, as described above. For each endogenous pre-mRNA examined in this study (PTC7, HPC2, or SRB2), gene-specific forward and reverse primers were designed to anneal to sequences in the two flanking exons. Following reverse transcription and PCR amplification, two products were expected: a larger product derived from the unspliced transcript containing the intron and a smaller product derived from the spliced transcript lacking the intron. The gene-specific primers were: HPC2 forward, 5’GACCATATTCAAACGATTGG3’, and HPC2 reverse, 5’AATTCTTCAGCGATGTTGGG3’; PTC7 forward, 5’TTTGCAAACGTTGGATTTAG3’, and PTC7 reverse, 5’GTCTTCTCCTGTAGGTGATC3’; SRB2 forward, 5’AAGCACAGTAGCAATCCATC3’, and SRB2 reverse, 5’ACGTTATCGAGTACATGAGC3’. Primers targeting U2 snRNA were used as an internal control: U2 forward, 5’ACGAATCTCTTTGCCTTTTGGC3’, and U2 reverse, 5’GTATTGTAACAAATTAAAAGG3’

### Primer extension assay

Splicing lariat intermediates and products were analyzed by primer extension. Total RNA was isolated, as described above. Typically, 8 μg of total RNA was resuspended in 5.4 μl ddH2O. 1.6 μl of annealing buffer (250 mM Tris-HCl, pH 8.3, 300 mM NaCl, 50 mM DTT) and 1 μl (100nM) of 5’-^32^P-labeled primer were added. The mixture was heated at 90℃ for 3min, transferred to 55℃ for 1min, then chilled on ice. A volume of 12 μl of reverse transcription (RT) mixture containing 18 mM Tris-HCl, pH 8.3, 21 mM NaCl, 3.6 mM DTT, 11 mM Mg(OAc)_2_, 0.72 mM dNTPs, and 0.25 unit of AMV reverse transcriptase (Promega), was added. The RT reaction was incubated at 42℃ for 20 min. The reaction was stopped by adding 350 μl G50 buffer (20mM Tris-HCl, pH7.5, 300mM sodium acetate, 2mM EDTA, and 0.25% sodium dodecyl sulfate), followed by PCA extraction and ethanol precipitation. The pellet was resuspended in ddH2O and 2×RNA loading dye (Fermentas) and heated at 95℃ for 4 min. Samples were loaded on 8% polyacrylamide gel (acrylamide/bisacrylamide 19:1) containing 8 M urea. Radioactive reverse transcription products were visualized by autoradiography. The primer 5’ GGCACTCATGACCTTC 3’ was complementary to *CUP1*, a 3’ exon sequence shared in common by pre-mRNA, mRNA, and lariat 2/3 intermediate. Exon-Exon junction primers were also used: 5’ GTACGTCGACCAGAATC 3’ complementary to Exon 1-Exon 2, and 5’ CCAACCTCAGAATCCATTGTTAATTC 3’ complementary to Exon 1-Exon 3. U6 primer 5’ CGGTTCATCCTTATGCAG 3’ and U2 primer 5’ GTATTGTAACAAATTAAAAGG 3’ were used as controls. Relative levels of reverse transcription stop bands were quantified by ImageJ (version 2.1.0).

### U2 pseudouridylation and Prp5 synthetic lethality assay

Yeast strain yYu (GB-53) (*MATa, ade2 cup1-Δ::ura3 leu2 lys2 trp1 prp5-Δ::loxP pus-7 Δ::HIS3 snR81-Δ::kanMX pus1-Δ::hphMX6, pRS316-PRP5[PRP5 URA3 CEN ARS]*) (Wu et al. 2016) was co-transformed with one of the pRS314-PRP5 variants (Prp5 WT, GAR, or N399D) (Wu et al. 2016) and one of the YEplac181 variants expressing designer guide RNAs directing U2 pseudouridylation at various positions within the BSRR or an empty vector. Transformants were selected on solid media lacking tryptophan and leucine. Colonies were picked and grown in selective media, and equal numbers of cells were plated on 5-FOA plates. Plates were incubated at 30°C for 4 days and photographed.

### Prp5-U2 and Prp5-U2-pre-mRNA Binding Assays

A derivative of yYZX02 (*pus7Δ, snR81Δ, pus1Δ*) expressing FLAG-tagged Prp5 from pRS314-PRP5-FLAG was generated as described previously (Wu et al. 2016). YEplac181 plasmids directing U2 pseudouridylation to generate the Ψ40Ψ42Ψ44Ψ45, Ψ38Ψ40Ψ42Ψ44Ψ45, and Ψ35Ψ40Ψ42Ψ44Ψ45 variants were introduced together with the *ACT1-CUP1* reporter plasmid (YEplac195, URA3) containing the weak BSS GACAGAC. Transformants were selected on SD-Leu/-Ura/-Trp medium and grown to an OD600 of 3.5 prior to extract preparation.

Briefly, cell pellets harvested from a 500 mL culture were resuspended in 25 mL of ice-cold Yeast cell lysis Buffer (50 mM Tris-HCl pH 7.5, 150 mM NaCl, 5 mM MgCl_2_, 1 mM EDTA, 10% glycerol, 1 mM DTT, and 1 mM PMSF) and lysed by two passes through a French Press (100-120 MPa). The cell lysate was clarified by centrifugation at 12,000 × g for 30 min at 4 °C to collect the supernatant.

For Co-IP, anti-FLAG beads (Thermo Scientific™ Pierce™ Anti-DYKDDDK Magnetic Agarose Beads) pre-washed with ice-cold lysis buffer were rotated with cell extracts at 4 °C for 4 h. The beads were then separated magnetically, washed three times with IP Wash Buffer (20 mM HEPES-KOH pH 7.9, 150 mM KCl, 1.5 mM MgCl2, 0.1% NP-40, 10% glycerol, 1 mM DTT, and 1 mM PMSF), and split into two portions for protein and RNA analyses.

For SDS-PAGE protein analysis, the washed beads were incubated in 2× SDS-PAGE Loading Buffer (4% SDS, 10% 2-mercaptoethanol, 20% glycerol, 0.125 M Tris-HCl pH 6.8, and 0.004% bromophenol blue) at 95 °C for 3 min, after which the supernatant was collected after brief centrifugation, loaded on the SDS-PAGE gel, and analyzed by western blotting using anti-FLAG antibody.

For RNA analysis, Phenol:Chloroform:Isoamyl alcohol (PCA, 25:24:1) was added directly to the washed beads and incubated at 70 °C for 10 min. After centrifugation, the aqueous phase was re-extracted with PCA, and RNA was precipitated with ethanol. U1, U2, and *ACT1-CUP1* pre-mRNA were quantified by RT-PCR using primers specific for each RNA species.

## Acknowledgements

We thank members of the Yu laboratory for extremely helpful discussions throughout this work. We also thank Dr. Charles Query for providing *PRP5* strains used in this study, and Drs. Eric Phizicky, Elizabeth Grayhack, and Xin Li for sharing reagents. This work was supported by National Institutes of Health grant R01GM138387 (to YTY).

## Author Contributions Statement

RZ and YTY conceived the study and wrote the manuscript. RZ and MDD performed most of the experiments, whereas JC, HA, and YS contributed to specific experiments. All authors approved the manuscript.

